# Temporal attention amplifies stimulus information in fronto-cingulate cortex at an intermediate processing stage

**DOI:** 10.1101/2024.03.06.583738

**Authors:** Jiating Zhu, Karen J. Tian, Marisa Carrasco, Rachel N. Denison

## Abstract

The human brain faces significant constraints in its ability to process every item in a sequence of stimuli. Voluntary temporal attention can selectively prioritize a task-relevant item over its temporal competitors to alleviate these constraints. However, it remains unclear when and where in the brain selective temporal attention modulates the visual representation of a prioritized item. Here, we manipulated temporal attention to successive stimuli in a two-target temporal cueing task, while controlling for temporal expectation with fully predictable stimulus timing. We used MEG and time-resolved decoding to track the spatiotemporal evolution of stimulus representations in human observers. We found that temporal attention enhanced the representation of the first target around 250 milliseconds after target onset, in a contiguous region spanning left frontal cortex and cingulate cortex. The results indicate that voluntary temporal attention recruits cortical regions beyond the ventral stream at an intermediate processing stage to amplify the representation of a target stimulus. This routing of stimulus information to anterior brain regions may provide protection from interference in visual cortex by a subsequent stimulus. Thus, voluntary temporal attention may have distinctive neural mechanisms to support specific demands of the sequential processing of stimuli.

**Significance statement:** When viewing a rapid sequence of visual input, the brain cannot fully process every item. Humans can attend to an item they know will be important to enhance its processing. However, how the brain selects one moment over others is little understood. We found that attending to visual information at a precise moment in time enhances visual representations around 250 ms after an item appears. Unexpectedly, this enhancement occurred not in the visual cortex, but in the left fronto-cingulate cortex. The involvement of frontal rather than posterior cortical regions in representing visual stimuli has not typically been observed for spatial or feature-based attention, suggesting that temporal attention may have specialized neural mechanisms to handle the distinctive demands of sequential processing.

## Introduction

Attention as a cognitive process allows us to select the most relevant sensory information to guide our behavior given our limited processing resources. Attentional selection happens not only across space but also across time as we process continuous visual input in our dynamic environment (***Carrasco, 2011; Anton-Erxleben and Carrasco, 2013; Denison et al***., ***2017; Nobre and Van Ede, 2023; Denison, 2024***). The goal-directed prioritization of a task-relevant time point is voluntary temporal attention (***Nobre and Van Ede, 2018; Denison et al., 2021***). For example, when returning a table tennis serve, we voluntarily attend to the ball at the moment it bounces on the table, because it is critical to see the ball at this time to successfully return the serve (***Land and Furneaux, 1997***). Attending earlier or later is less useful for predicting the trajectory of the ball.

In the temporal domain, limitations in continuous visual processing are often studied by using a rapid sequence of stimuli, in which observers are asked to prioritize one or more events. Various behavioral findings indicate that the brain cannot fully process each stimulus in a rapid sequence (***Lawrence, 1971; Raymond et al., 1992***). In the attentional blink, detection accuracy for the second of two target stimuli suffers when the stimuli are separated by 200-500 ms (***Raymond et al., 1992; Dux and Marois, 2009***). In temporal crowding, the identification of a target stimulus is impaired when it is surrounded by other stimuli in time, across similar intervals of 150-450 ms (***Yeshurun et al., 2015; Tkacz-Domb and Yeshurun, 2021***). At this timescale, voluntary temporal attention can flexibly prioritize stimuli at relevant time points, improving perceptual sensitivity and reaction time for temporally attended stimuli at the expense of the processing of stimuli earlier and later in time, effectively reducing temporal constraints by selecting one stimulus over others (***Denison et al., 2017, 2021; Fernández et al., 2019; Duyar et al., 2023, 2024***).

Despite the behavioral evidence for selectivity in temporal attention, little is known about the neural mechanisms underlying the ability to selectively attend to one point in time over another. Neural correlates of temporal anticipation have generally been studied by manipulating the timing predictability of a single target stimulus (***Coull and Nobre, 1998; Correa et al., 2006; Anderson and Sheinberg, 2008; Van Ede et al., 2018; Lima et al., 2011; Nobre and Van Ede, 2018, 2023***). Predictability increases the firing rates of inferotemporal neurons in non-human primates (***Anderson and Sheinberg, 2008***) and the amplitude of visual evoked potentials in human EEG around 100-150 ms after stimulus onset (***Doherty et al., 2005; Correa et al., 2006***), and induces anticipatory activity in human EEG before the expected time (***Samaha et al., 2015; Breska and Ivry, 2020***). However, studies with a single target stimulus cannot disentangle the process of attending to a task relevant time point from processes associated with the temporal predictability of the target onset, or temporal expectation. In the spatial and feature-based domains, attention and expectation can have distinct behavioral and neural effects, indicating the importance of experimentally dissociating these two processes (***Summerfield and Egner, 2009; Wyart et al., 2012; Summerfield and Egner, 2016; Rungratsameetaweemana and Serences, 2019; Moerel et al., 2022***). In addition, mechanisms for selecting a task-relevant stimulus from a sequence may differ from those involved in enhancing a single stimulus with no other temporally proximal stimuli. This is because multiple stimuli in a rapid sequence may create competition for processing resources that a single, isolated stimulus does not.

Consequently, it remains unknown how humans use voluntary temporal attention to flexibly select a relevant stimulus representation within a sequence, at the expense of temporal competitors. Specifically, it is unclear what stage or stages of visual processing are affected by temporal attention. Temporal attention, like spatial attention (***van Es et al., 2018; Dugué et al., 2020; Liu et al., 2021***) and feature-based attention (***Maunsell and Treue, 2006; Liu et al., 2007; Foster and Ling, 2022***) could affect early visual representations, and there is initial evidence that temporal attention can improve the reliability of visual responses (***Denison et al., 2024***). Alternatively or in addition, temporal attention could affect later visual representations or the transfer of stimulus information to downstream processing stages.

To study how temporal attention meditates selection, we therefore designed a minimal stimulus sequence with two temporally predictable stimuli on each trial (***Denison et al., 2017, 2021***). Only the time point to be attended, indicated at the beginning of each trial by a precue, varied across trials. With timing predictability controlled, differences in the neural representations of a stimulus when it was temporally attended vs. unattended could be attributed to temporal attentional selection. We used MEG together with this psychophysical task to investigate when and where in the brain selective temporal attention affects representations of visual stimuli. Our behavioral results confirmed that temporal attention improved perceptual sensitivity and speeded reaction time.

Using time-resolved decoding, we found that voluntary temporal attention enhanced the orientation representation of the first grating target 235-300 ms after target onset, an intermediate time window following the earliest visual evoked responses. This time interval is consistent with temporal processing constraints revealed behaviorally by tradeoffs due to voluntary temporal attention (***Denison et al., 2017, 2021***), the attentional blink (***Raymond et al., 1992***), and temporal crowding (***Yeshurun et al., 2015***). In source space reconstructions, we found that although orientation decoding was strongest in occipital areas, as expected, the strongest effects of temporal attention on orientation representations appeared in left fronto-cingulate regions. Additionally, we found no impact of temporal attention on univariate visual responses, unlike previous studies that manipulated temporal attention without isolating its effect from temporal expectation. Altogether the results suggest that voluntary temporal attention selectively prioritizes a target stimulus by amplifying its representation in fronto-cingulate regions at an intermediate processing stage around 250 ms, perhaps to protect it from a subsequent temporal competitor in visual cortex. This result suggests that temporal attention achieves stimulus selection using neural mechanisms for routing a stimulus representation not typically observed for spatial or feature-based attention, perhaps due to the distinctive demands of sequential processing.

## Results

### Temporal precueing improved perceptual sensitivity

To investigate the effects of voluntary temporal attention, we recorded MEG while observers performed a two-target temporal cueing task (Figure 1A). At the start of each trial, a precue tone instructed observers to attend to either the first target (T1) or the second target (T2). The two sequential grating targets were separated by a 300 ms stimulus onset asynchrony (SOA). At the end of each trial, a response cue tone instructed observers to report the tilt (clockwise or counterclockwise) of one of the targets. The precue and response cue were matched on 75% of the trials (valid trials) and mismatched on 25% of the trials (invalid trials), so observers had an incentive to direct their attention to the precued target.

**Figure 1.**
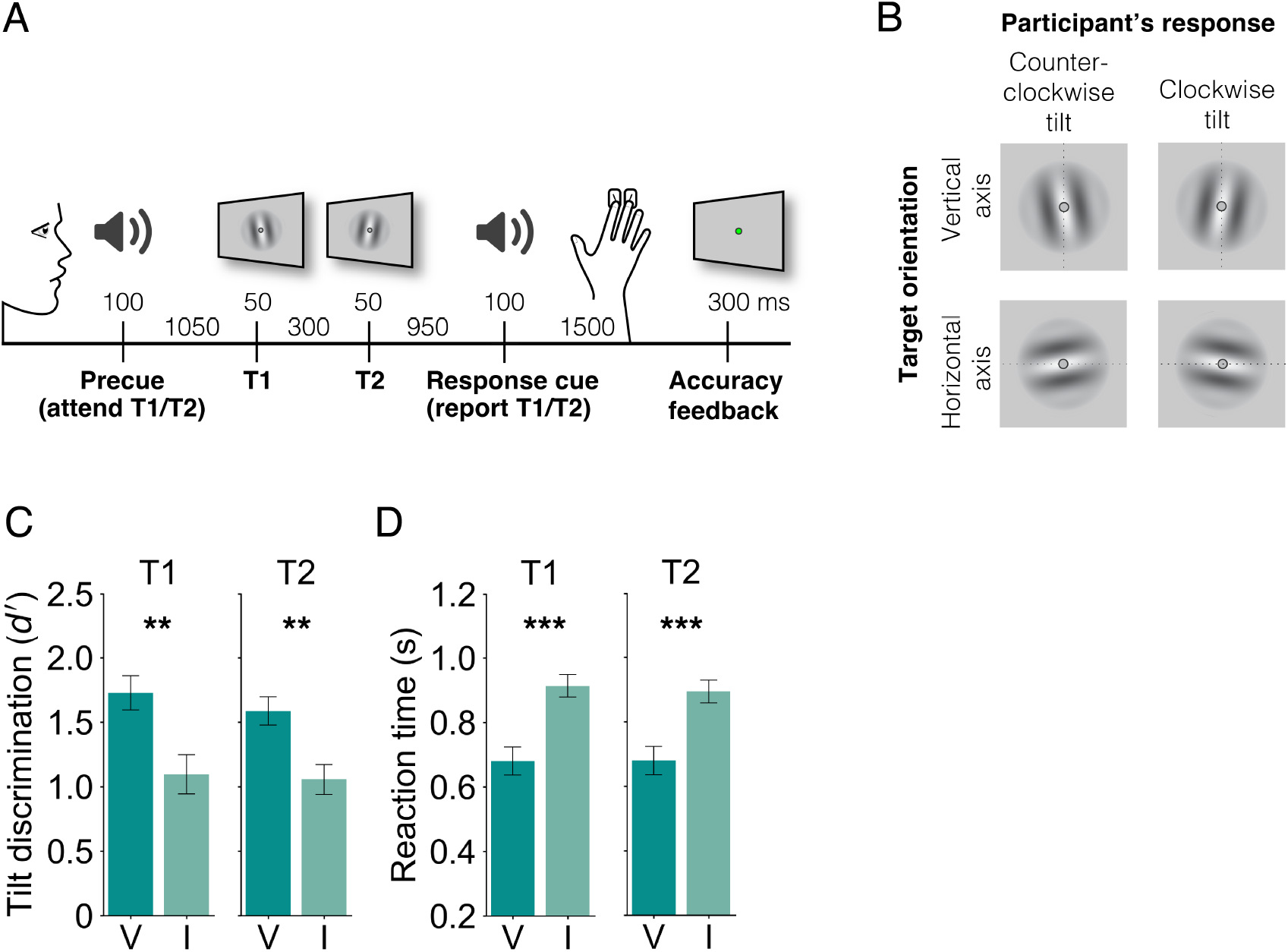
Two-target temporal cueing task and behavioral results. (A) Trial timeline showing stimulus durations and SOAs. Precues and response cues were pure tones (high = cue T1, low = cue T2). (B) Targets were independently tilted about the vertical or horizontal axes. Participants were instructed to report the clockwise or counterclockwise tilt of the target indicated by the response cue, and axis orientation was the decoded stimulus property. (C) Tilt discrimination (sensitivity) and (D) reaction time for each target (T1, T2) by validity condition. Sensitivity was higher and reaction time was faster for valid (V) than invalid (I) trials. Error bars indicate ±1 SEM. ** p *<* 0.01; *** p *<* 0.001.

Importantly, targets were tilted independently about either the vertical or the horizontal axis, allowing us to use MEG to decode a sensory feature — axis orientation — that was orthogonal to the participant’s report. Targets were oriented near vertical or horizontal, with individually titrated tilt thresholds ranging from 0.4-1.5 degrees (mean 0.76 degrees), and the participant’s report was clockwise or counterclockwise tilt with respect to the main axis (Figure 1B).

Temporal attention improved tilt discrimination performance, consistent with previous findings (***Denison et al., 2017, 2021; Fernández et al., 2019; Rohenkohl et al., 2014; Samaha et al., 2015; Duyar et al., 2024***). Perceptual sensitivity (*d*^’^) was higher for valid trials than invalid trials (Figure 1C; main effect of validity: 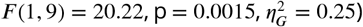 Perceptual sensitivity was similar for targets T1 and T2. The improvement in *d*^’^ with temporal attention was significant for both target 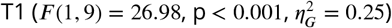 and target 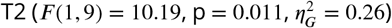 There was no main effect of target or interaction between validity and target (*F* (1, 9) *<* 0.59, p *>* 0.47).

Reaction time (RT) was faster for valid than invalid trials (Figure 1D; main effect of validity: 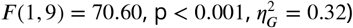 with improvements for both target 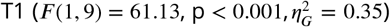 and target 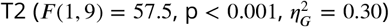. There was no main effect of target or interaction between validity and target (*F* (1, 9) *<* 0.67, p *>* 0.43). Therefore the improvement in perceptual sensitivity with the precue was not due to a speed-accuracy tradeoff.

### No effect of temporal attention on visual evoked response peaks

We first investigated whether temporal attention affected univariate visual evoked responses recorded from MEG. To do so, we identified visually responsive channels for each participant and session by ranking all 157 channels by the magnitudes of their visually evoked responses following stimulus onset, regardless of the precue (see Methods). The average for the 5 most visually responsive channels, which were in posterior locations, shows clear visual evoked responses after each target (Figure 2A), with no apparent effect of temporal attention. We characterized the evoked response peaks quantitatively and found no differences between precue conditions in the peak amplitudes of the evoked responses for any selected number of channels (Figure 2B; T1: *F* (1, 9) *<* 1.31, p *>* 0.28; T2: *F* (1, 9) *<* 5.2 and p *>* 0.043 uncorrected; none survived corrections for multiple comparisons across channel groupings). Likewise, we observed no differences in evoked response peak latencies (Figure 2C; T1: *F* (1, 9) *<* 1.67, p *>* 0.23; T2: *F* (1, 9) *<* 2.65, p *>* 0.14). Thus we found no evidence that voluntary temporal attention affected visual evoked responses, when assessed in a univariate fashion.

**Figure 2.**
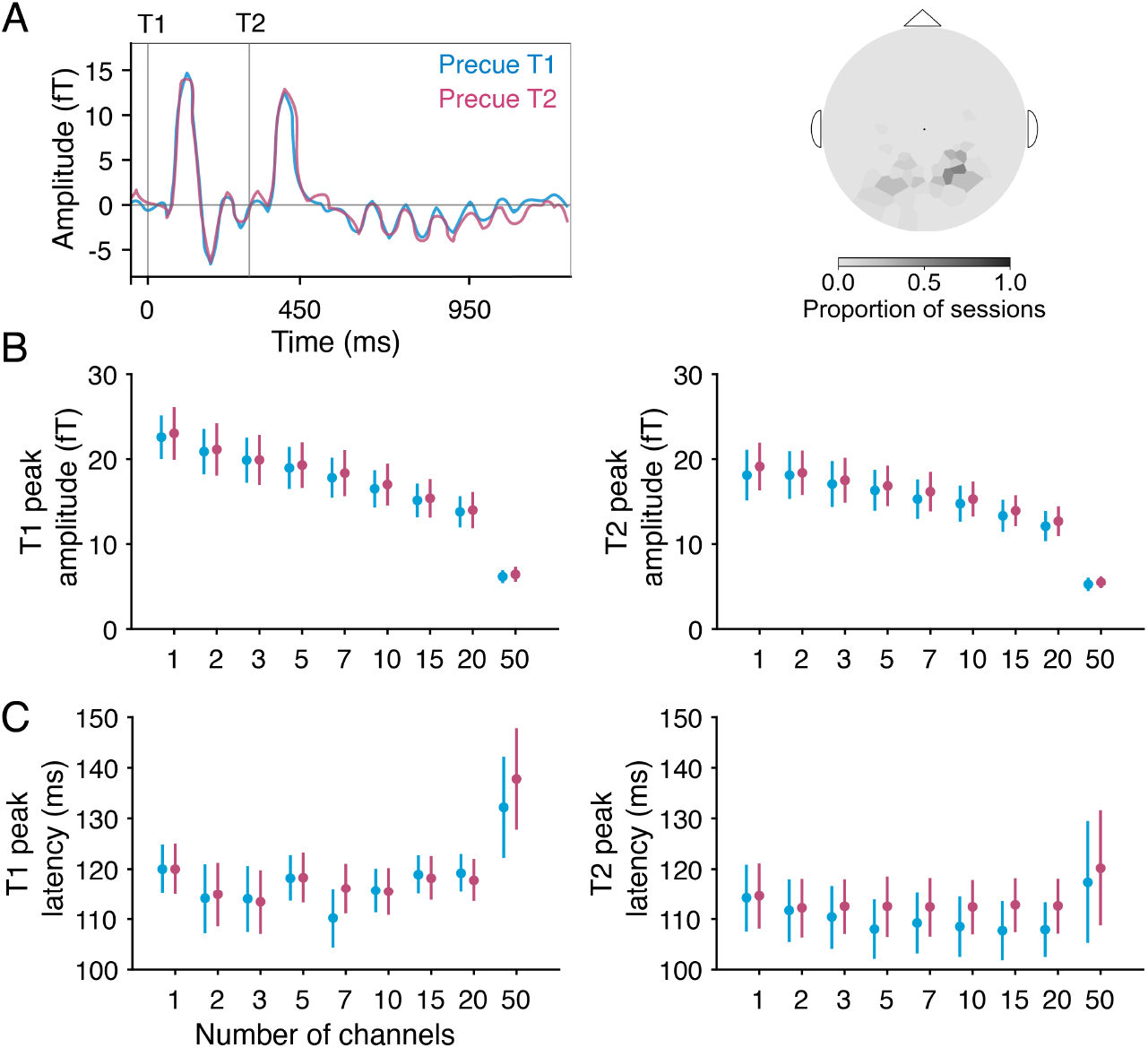
MEG evoked responses. (A) Average evoked time series by precue from the 5 most visually responsive channels. Channels were rank ordered by evoked peak prominence. Target onsets are marked with gray vertical lines. Varying the number of selected channels yielded no differences in target-evoked (B) peak amplitude or (C) peak latency by precue for either target in any channel grouping.

### Temporal attention increased orientation decoding performance following the initial visual evoked response

To investigate whether temporal attention improved the representation of orientation information, we next examined multivariate patterns from the MEG channels, using decoding accuracy as an index of the quality of orientation representations. For each participant and session, we selected the 50 most visually responsive channels for decoding analysis (see Methods). As expected, the selected channels tended to be in posterior locations (Figure 3 inset at top right).

**Figure 3.**
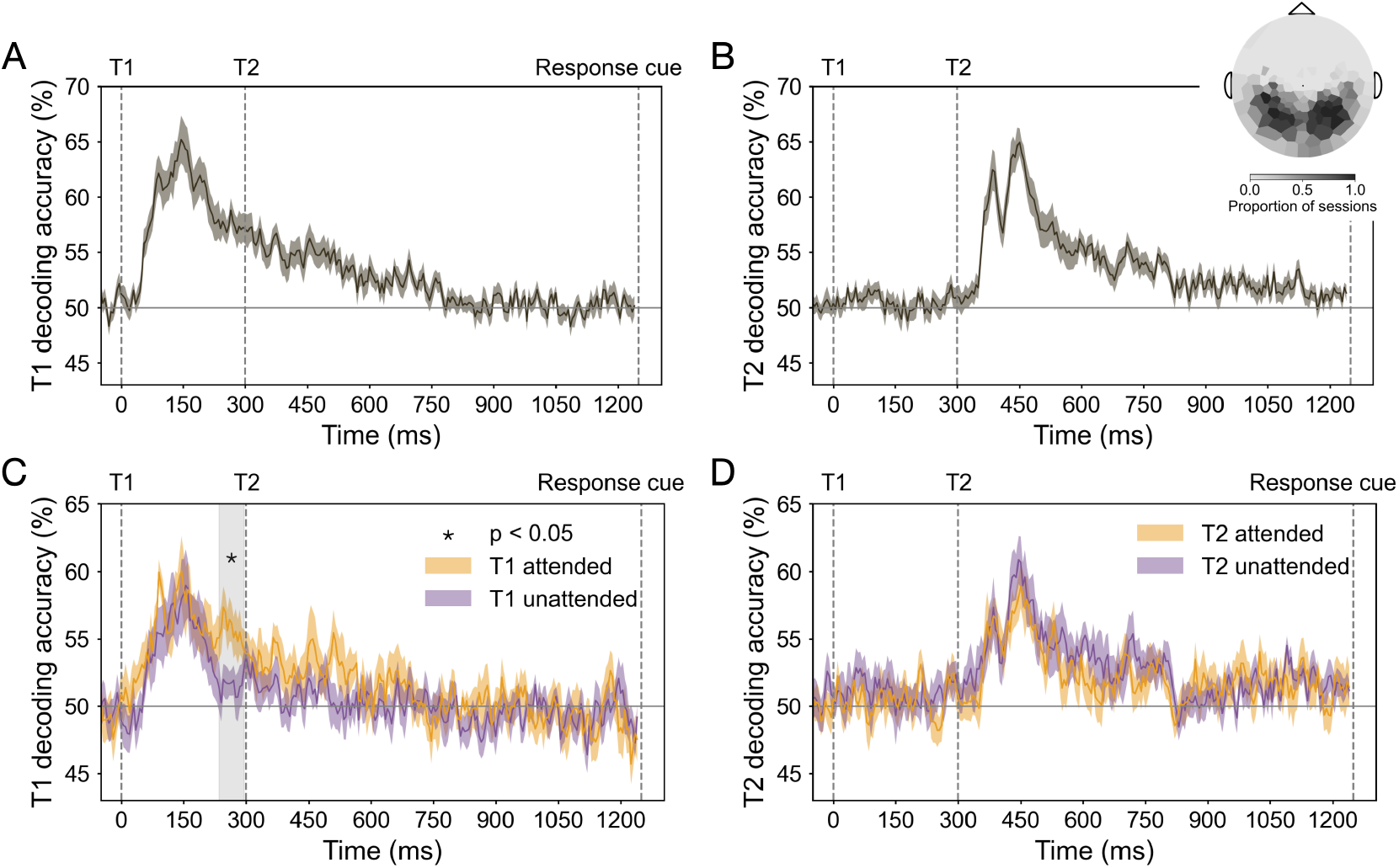
Decoding performance in MEG sensor space. Event onsets are marked with vertical dashed lines. (A) T1 orientation decoding performance for all trials. (B) T2 orientation decoding performance for all trials. Inset in (B) shows the topography of channels used for decoding across all sessions (the 50 most visually responsive channels per session). (C) T1 orientation decoding performance for target attended (precue T1) and unattended (precue T2) trials. Enhancement of orientation representation occurred 235-300 ms after target onset; gray shaded region shows cluster-corrected significant window (“critical time window”). (D) T2 orientation decoding performance for target attended (precue T2) and unattended (precue T1) trials.

We trained separate orientation classifiers for T1 and T2, which on each trial had independent vertical or horizontal axis orientations. For both targets, decoding performance reached about 65% accuracy, peaking around 150 ms after target onset (Figure 3A and B). There was no significant difference between the peak decoding performance of the two targets (decoding accuracy at 150 ±25 ms, t = 1.81, p *>* 0.10). Therefore, stimulus orientation was decodable for both targets, with comparable performance for T1 and T2, allowing us to investigate the time-resolved orientation representation of each target separately. Note this method cannot be used to decode the stimulus orientation before target onset, because according to our task design, no orientation information is available before the target appears. Thus this analysis would not be able to identify prestimulus effects of temporal attention vs. temporal expectation.

To investigate the effect of temporal attention on the orientation representation of each target, we next trained and tested time-resolved classifiers on target attended trials and unattended trials separately. T1 decoding accuracy was higher on attended than unattended trials in a time window 235-300 ms after target onset (p *<* 0.05 cluster-corrected; Figure 3C). This significant window started about 100 ms after orientation decoding performance peaked and ended just before T2 appeared. There was no similar enhancement when decoding the T2 orientation (Figure 3D).

We confirmed the enhancement of temporal attention on orientation decoding for T1 around 250 ms in a separate dataset, in which the targets were superimposed on a 20-Hz flickering noise patch instead of a blank background (see Supplementary Figure 2 and Supplementary Text). Again we found no significant attentional enhancement for T2. The enhancement of the orientation representation for T1 around 250 ms in two datasets confirms the robustness of this finding and its specificity to the first target.

### Widespread decodability of orientation representations across cortex

We next asked how orientation representations and the effect of temporal attention varied across the cortex. We focused our spatial analysis on T1, because we found no effect of temporal attention on T2 decoding at the channel level. Using source reconstruction, we estimated the MEG response at each time point at vertex locations across the cortical surface (***Dale et al., 2000; Gramfort et al., 2013***). We then applied time-resolved decoding analysis to vertices in each of 34 bilateral Desikan-Killiany (DK) atlas regions of interest (ROIs) (***Desikan et al., 2006***). MEG source reconstruction has spatial specificity on the order of *<*5 cm for cortical brain areas (***Hauk et al., 2011; Hedrich et al., 2017; Gross, 2019***), and the spatial scale of the DK atlas is typical for reporting MEG decoding performance (***van de Nieuwenhuijzen et al., 2013; King et al., 2016; Ramkumar et al., 2016; de Vries et al., 2021***). To be conservative, we report all main findings at the scale of cortical lobes. In the critical time window, orientation decoding performance across all trials was highest in posterior regions, as expected (Figure 4A). We obtained decoding performance for the occipital, parietal, temporal, and frontal lobes by averaging decoding performance across the ROIs within each lobe (***Klein and Tourville, 2012***).

**Figure 4.**
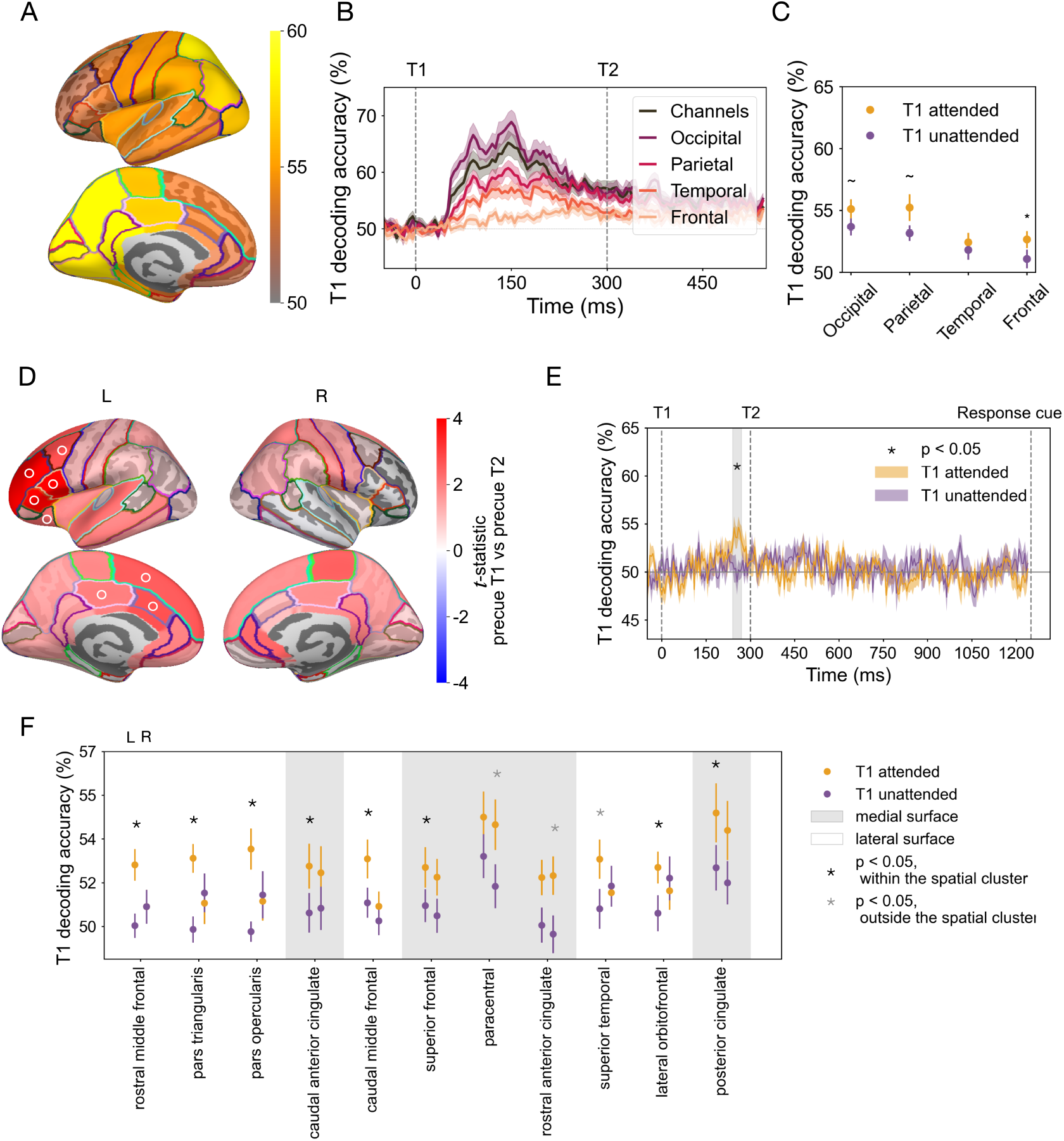
Decoding performance in MEG source space and topography of temporal attentional enhancement of orientation representations. (A) T1 decoding performance for 34 bilateral Desikan–Killiany atlas regions averaged across time points within the critical time window. The regions with the highest decoding performance were posterior regions in light yellow. (B) T1 decoding performance from all trials by lobe. Consistent with T1 decoding performance from sensor space, decoding performance for the occipital, parietal, and temporal lobes peaked around 150 ms after target onset, whereas frontal decoding peaked later, around 250 ms. (C) Effect of temporal attention averaged across time points within the critical time window. Error bars indicate ±1 SEM. * p *<* 0.05, ∼ p *<* 0.1. (D) T1 decoding differences between attended and unattended conditions for left (L) and right (R) hemispheres, based on the average decoding performance within the critical time window for each of 68 DK ROIs. A connected left fronto-cingulate region survived spatial cluster correction (p *<* 0.05, ROIs in the cluster marked with ○ symbol). (E) Time-resolved decoding accuracy of the cluster (average across the 8 ROIs marked with ○ in (D)) recovers enhancement of orientation representation within the critical time window (240-275 ms after target onset, gray shaded region). (F) Left-lateralization of effect of temporal attention on T1 decoding in the critical time window. ROIs ordered by their attended vs. unattended p-values, based on the hemisphere with the strongest attention effect (all ROIs with uncorrected p *<* 0.05 in at least one hemisphere are shown). ROIs on the lateral surface of the cortex (white background) show strong left lateralization of temporal attention. ROIs on the medial surface (gray background) show more bilateral effects of temporal attention. Error bars indicate ±1 SEM.

The T1 decoding performance for each of the 4 lobes at each time point showed a systematic pattern of decoding accuracy: highest in occipital, lower in parietal and temporal, and lowest in the frontal lobe (Figure 4B). In addition, decoding performance peaked later in the frontal lobe than in the other three lobes, around 250 ms. Such progression of decoding strength and timing across lobes is consistent with the visual processing hierarchy, demonstrating the feasibility of decoding orientation in source space.

### Temporal attention enhanced orientation representations in left fronto-cingulate cortex

We next asked where in the brain temporal attention increases orientation representations of T1 during the critical time window (235-300 ms after target onset; Figure 3). In this time window, although the frontal lobe had lower decoding overall, it showed the biggest difference between attended and unattended trials (Figure 4C), which was statistically reliable 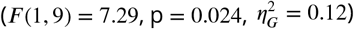 The occipital lobe 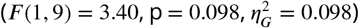 and the parietal lobe 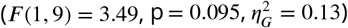 showed marginal differences between attention conditions, while the temporal lobe had no statistically significant difference (*F* (1, 9) *<* 0.39, p *>* 0.54).

To more precisely localize the cortical regions underlying the enhancement of orientation representations, we examined orientation decoding in the 34 DK ROIs in each hemisphere within the critical time window (235-300 ms) in which temporal attention improved T1 decoding in sensor space. One spatial cluster showed an attentional enhancement of orientation decoding that survived the cluster permutation correction across ROIs (regions in the cluster are marked by ○ in Figure 4D). This significant cluster was comprised of eight regions in the left hemisphere: seven in the frontal lobe and one in the parietal lobe. The eight regions ranked by their p-values are left rostral middle frontal, pars triangularis, pars opercularis, caudal anterior cingulate, caudal middle frontal, superior frontal, lateral oribital frontal, and posterior cingulate. If we treat the cingulate cortex as a separate lobe (***Klein and Tourville, 2012***), two of the regions in the cluster, including the parietal region, are in the cingulate lobe. Therefore we characterize the significant cluster as located in left fronto-cingulate cortex.

We next investigated the time-resolved decoding performance in the fronto-cingulate cluster by averaging across the eight regions at every time point. Orientation decoding performance was enhanced in target attended trials in a time window (240-270 ms) which fell within the significant time window we found in the sensor space (Figure 4E), confirming that the critical time window was recovered from the fronto-cingulate cluster alone. In addition, decoding performance in the cluster peaked around 250 ms in target attended trials, with no transient early peak as was found in the occipital lobe. This time course indicates that the orientation information decoded from the fronto-cingulate cluster did not arise from signal leakage from the occipital lobe during source reconstruction.

Finally, we investigated the degree of hemispheric lateralization in the regions with the strongest attention effects (Figure 4F). Regions located on the lateral surface of the hemisphere were strongly lateralized, with significant differences between attended and unattended trials for regions in the left hemisphere but not in their right hemisphere counterparts, whereas medial regions tended to have bilateral attention effects. It is important to note that source estimation may not be sufficiently precise to fully localize signals arising from the medial surface to the correct hemisphere, due to the spatial proximity of the two hemispheres at the midline (***Molins et al., 2008***). At the same time, the bilateral pattern for these midline regions increases confidence that the signals originate from cingulate cortex rather than from more lateral frontal areas. Altogether, the source analysis reveals that the strongest temporal attentional enhancement of orientation representations was left-lateralized in the fronto-cingulate cortex.

## Discussion

The visual system faces significant constraints in processing the continuous visual information it receives. Humans can cognitively manage these constraints by using voluntary temporal attention to prioritize stimuli at task-relevant times at the expense of processing temporal competitors, but the neural mechanisms underlying this ability have received scant investigation. Here we experimentally manipulated temporal attention—while controlling temporal expectation—and used time-resolved MEG decoding (***Cichy et al., 2015; King et al., 2016***) together with source localization to uncover how voluntary temporal attention selectively enhances neural representations of oriented stimuli at task-relevant points in time within a stimulus sequence. Our results reveal neural mechanisms of temporal attentional selection, and, unexpectedly, argue for a specific role of the left fronto-cingulate cortex in amplifying target information under temporal constraints.

We found, in two independent datasets, that temporal attention enhanced the orientation representation of the first target at an intermediate processing stage around 250 ms: later than early visual event-related responses and the peak orientation decoding accuracy (∼120-150 ms after target onset) (***Cichy et al., 2015; Wardle et al., 2016; Pantazis et al., 2018***), but before decoding performance fell to chance (∼500 ms), and just before the onset of T2. Interestingly, this window corresponds to the interval when temporal attention is maximally selective. In ***Denison et al***. (***2021***), maximal attentional tradeoffs in behavior appeared when the two targets were separated by an SOA of 250 ms, with decreasing tradeoffs at shorter and longer SOAs.

Attention enhanced orientation decodability for T1 but not for T2, despite equivalent behavioral effects of temporal attention for both targets. This dissociation suggests that temporal attention modulates the processing of the two targets by different mechanisms. Such differences may not be surprising. Unlike in spatial attention, where attending to a stimulus on the left is not fundamentally different from attending to one on the right, temporal attention interacts with the directionality of time. T1 processing is not complete when T2 appears, while T2 appears in the context of ongoing T1 processing. One possibility is that temporal attention may route T1 information to the fronto-cingulate cortex to protect it from interference in the visual cortex, at a sensory encoding stage, when T2 appears. T2 information, on the other hand, does not need to be transferred to fronto-cingulate cortex, as there is no subsequent temporal competitor. Selection of T2 may take place instead at a later stage (e.g., during a decision readout). If attentional selection only affects a later stage of T2 processing, we would not be able to observe it with our current measure, which reflects a sensory feature of the stimulus that is orthogonal to the task-relevant stimulus information. The finding that attentional enhancement of orientation information was specific to the first target is consistent with a previous two-target temporal cueing study, which found that temporal attention increased the reliability of T1 responses (***Denison et al., 2024***). Thus, modulating the first target may be sufficient to bias downstream competition for processing T1 vs. T2. Previous behavioral results from a temporal attention task with three sequential targets have also suggested temporal asymmetries (***Denison et al., 2017***). Such specialized mechanisms for temporal attentional selection may reflect the demands of dynamic, sequential processing.

Although studies of temporal expectation have found modulations of visual cortical responses (***Doherty et al., 2005; Correa et al., 2006; Lima et al., 2011; Anderson and Sheinberg, 2008; Van Ede et al., 2018***), we found that the most reliable modulations of sensory representations by temporal attention were not in the ventral stream. Rather, the left frontal cortex and cingulate regions showed the strongest attentional modulations of orientation decoding, even though they had lower overall orientation decoding levels than occipital regions. Few studies have investigated how attention affects stimulus representations in frontal or cingulate cortices. Two studies reported better decoding of stimulus representations in precentral sulcus when the decoded feature dimension was task-relevant vs. irrelevant (***Ester et al., 2016; Yu and Shim, 2017***). Fronto-cingulate neurons also carry information about the location of spatial attention (***Kaping et al., 2011; Voloh et al., 2015; Westendorff et al., 2016***). Previous studies could not have uncovered effects of temporal attention on neural representations beyond visual areas, because they used electrode penetrations confined to sensory areas, or EEG methods which did not permit high spatial resolution source reconstruction. Taking advantage of the combined temporal and spatial resolution of MEG, the present results revealed which cortical areas were modulated by temporal attention during the precise time window when this modulation occurred.

The strong left lateralization we observed in frontal and cingulate areas is consistent with studies that have recorded a left hemisphere bias for temporal cueing using positron emission tomography (PET) and fMRI (***Nobre and Rohenkohl, 2014; Coull and Nobre, 1998; Coull et al., 2000, 2001; Davranche et al., 2011; Cotti et al., 2011***). In particular, the left inferior frontal gyrus (BA44/6) found in temporal orienting of attention (***Coull and Nobre, 1998***) overlaps with the pars opercularis region, which is one of the frontoparietal regions we found to have the strongest temporal attention effect. In these previous studies, univariate measures showed activity in these regions, but their precise function was unclear. One interpretation was that these frontoparietal regions could be part of a control network for the deployment of attention at specific time points (***Kastner and Ungerleider, 2000; Wang et al., 2010***). The current findings that these areas carry orientation-specific information, which is enhanced when temporally attended, suggest the alternative possibility that these areas are involved in maintaining attended stimulus representations. It is also possible that a left frontoparietal network is recruited for multiple aspects of temporal attention, including both control and stimulus selection.

The eight connected fronto-cingulate regions showing higher decoding performance for attended targets overlap substantially with regions that have been associated with the cingulo-opercular (CO) network (***Dworetsky et al., 2021***). The CO regions—the dorsal anterior cingulate cortex/medial superior frontal cortex (dACC/msFC) and anterior insula/frontal operculum (aI/fO)—show activity in diverse tasks (***Dosenbach et al., 2007***). In a visual working memory task, a retrocue directing focus to an item already in memory recruited the CO network (***Wallis et al., 2015***), suggesting that CO regions were selecting the cued item and reformatting it into an action-oriented representation (***Myers et al., 2017***). The CO network has also been found to flexibly affiliate with other networks depending on task demands in cognitive tasks with different combinations of logic, sensory, and motor rules (***Cocuzza et al., 2020***). Based on these findings, we might speculate that the CO network provides extra cortical resources to maintain and possibly reformat the representation of the first target, which might otherwise get overwritten by the second target within the visual cortex.

The possibility that the CO regions select the cued item and encode the reformatted representation could suggest that temporal attention contributes to prioritization in working memory by selecting stimulus information for working memory encoding or reformatting. However, it is unlikely that the transient enhancement of orientation representations we observed is tied to working memory maintenance. In typical working memory tasks, the delay period is several seconds (***Harrison and Tong, 2009; Sala and Courtney, 2007***), whereas here we found enhancement around 250 ms and indeed no above-chance orientation decoding after 800 ms.

Previous research supports the idea that temporal anticipation can protect target processing from a subsequent distractor. One study used a warning signal on some trials to cue observers to an upcoming target that could be followed after 150 ms by a distractor. When the warning signal was present, orientation decoding for the target was enhanced ∼200-250 ms after target onset (***Van Ede et al., 2018***), but only when the distractor was present, suggesting that the warning signal served to reduce distractor interference. Another study, on working memory, presented distractors at a predictable time during the retention interval, 1.1 s following the final memory target. Occipital alpha power and phase locking increased just before the distractor appeared and were associated with reduced impact of the distractor on memory performance (***Bonnefond and Jensen, 2012***). These studies, which involved different task types, temporal scales between targets and distractors, and measured neural signals, suggest that the brain may have diverse mechanisms for shielding target processing from temporally anticipated distractors. Here we isolated the contribution of voluntary temporal attention to enhancing target processing in the presence of temporal distractors, while controlling other voluntary and involuntary processes related to stimulus predictability and alerting, to reveal the flexible, top-down mechanisms of temporal selection. In this case, attention enhanced target stimulus representations even before the temporal competitor appeared.

Isolating temporal attention from other processes also indicated that the mechanisms of temporal attention may be distinct from those of temporal expectation. Studies that manipulated temporal attention together with temporal expectation by manipulating the timing predictability of a single target stimulus found enhancements in early visual evoked responses (***Doherty et al., 2005; Correa et al., 2006; Griffin et al., 2002; Anderson and Sheinberg, 2008***), which we did not observe in response to our targeted manipulation of temporal attention. Although it is difficult to reach a strong conclusion from the absence of an effect, we did observe attention-related changes in neural activity at intermediate time windows—confirming the sensitivity of our measurements—and found no evidence for effects of temporal attention on univariate evoked responses across a range of channel selections. It is therefore possible that previous observations of early modulations of visual responses were more closely linked to timing predictability than to the prioritization of a task-relevant time point per se.

Temporal attention may also affect early sensory processing in some other way than increasing visual evoked responses. A recent study from our group measured occipital cortical responses to a steady-state flickering stimulus with overlaid targets (***Denison et al., 2024***). Temporal attention to the first target transiently increased the effect of the target on the steady-state response ∼150 ms after target onset, demonstrating early modulations specific to temporal attention. In our current data, we also observed an early peak in decoding accuracy for T1 that was present when T1 was attended but absent when it was unattended, which was localized to occipital and parietal regions. However, this difference between attention conditions did not survive cluster correction across the whole time series (see Supplementary Figure 3 and Supplementary Text), likely due to the brief duration of the peak.

Indeed, here when isolating temporal attention from temporal expectation, we found the strongest effects of temporal attention in fronto-cingulate cortex. Temporal cueing studies that combined temporal attention and expectation were not able to investigate stimulus representations in these anterior brain regions. Therefore, although the present results suggest distinct mechanisms for temporal attention and temporal expectation, future studies that independently manipulate these two processes in the same experiment will be important for resolving their shared and distinct mechanisms.

## Conclusions

We found that using voluntary temporal attention to select one stimulus over another within a short sequence enhanced the neural representation of the selected stimulus identity. This enhancement occurred around 250 milliseconds after the onset of the first target, reflecting an intermediate stage of processing that matches the timing of maximal temporal attentional tradeoffs observed behaviorally (***Denison et al., 2021***). Surprisingly, the enhancement was localized not to visual cortical regions but to left-lateralized fronto-cingulate cortex. The results suggest that temporal attention improves visual task performance by routing target information to these anterior regions, which may act as a protective reservoir for task-relevant information in the presence of a subsequent temporal competitor. In contrast, we found no effect of temporal attention—when isolated from temporal expectation—on visual evoked responses. The results thus revealed a role for cortical areas beyond the ventral stream in the temporal selection of a behaviorally relevant target and uncovered an unforeseen effect of voluntary temporal attention.

## Methods

### Observers

Ten observers (5 females, mean age = 29 years old, SD = 4 years), including authors RND and KJT, participated in the study. Each observer completed 1 behavioral training session and two 2-hour MEG sessions on separate days for 20 sessions of MEG data in total. This approach allowed us to check the reliability of the data across sessions for each observer and is similar to the approach taken by other MEG studies of vision (***Kok et al., 2017; Besserve et al., 2007***). All observers had normal or corrected-to-normal vision using MR safe lenses. All observers provided informed consent and were compensated for their time. Experimental protocols were approved by the University Committee on Activities involving Human Subjects at New York University.

### Stimuli

Stimuli were generated using MATLAB and Psychtoolbox (***Brainard and Vision, 1997; Pelli and Vision, 1997; Kleiner et al., 2007***) on an iMac.

Stimuli were projected using a InFocus LP850 projector (Texas Instruments, Warren, NJ) via a mirror onto a translucent screen. The screen had a resolution of 1024 × 768 pixels and a refresh rate of 60 Hz and was placed at a viewing distance of 42 cm. Stimuli were displayed on a medium gray background with a luminance of 206 cd/m^2^. Target timing was checked with photodiode measurements. For behavioral training sessions outside of the MEG, stimuli were presented on a gamma-corrected Sony Trinitron G520 CRT monitor with a resolution of 1024 × 768 pixels and a refresh rate of 60 Hz placed at a viewing distance of 56 cm. Observers were seated at a chin-and-head rest to stabilize their head position and viewing distance.

### Visual targets

Visual targets were full contrast sinusoidal gratings with spatial frequency of 1.5 cpd presented foveally. The gratings were 4° in diameter and had an outer edge subtending 0.4° that smoothly ramped down to zero contrast.

### Auditory cues

Auditory precues and response cues were pure sine wave tones 100 ms in duration with 10 ms cosine amplitude ramps at the beginning and end to prevent clicks. Tones were either high-pitched (1046.5 Hz, C6) indicating T1 or low-pitched (440 Hz, A4) indicating T2.

### Task

Observers were asked to direct voluntary temporal attention to different time points in a sequence of two visual targets and to discriminate the tilt of one target. On each trial, two targets (T1 and T2) appeared one after another in the same location. The targets were presented for 50 ms each and separated by a 300 ms stimulus onset asynchrony (SOA) based on psychophysical studies that have shown temporal attentional tradeoffs at this timescale (***Denison et al., 2017, 2021; Fernández et al., 2019***). Each target was tilted slightly clockwise (CW) or counterclockwise (CCW) from either the vertical or horizontal axis (Figure 1B). Tilts and axes were independent and counterbalanced for each target.

An auditory precue 1,050 ms before the targets instructed observers to attend to either T1 (high tone) or T2 (low tone). An auditory response cue 950 ms after the targets instructed observers to report the tilt (CW or CCW) of either T1 or T2. Observers pressed one of two buttons to indicate whether the tilt was CW or CCW relative to the main axis within a 1500 ms response window. At the end of the trial, observers received feedback for their tilt report via a color change in the fixation circle (green: correct; red: incorrect; blue: response timeout).

On every trial, the targets were fully predictable in time following the precue. The attended target varied trial-to-trial according to the precue, and the target selected for report varied trial-to-trial according to the response cue. On trials in which the precue directed attention to one target (80% of trials), the precue and response cue usually matched (75% validity), so the observers had an incentive to direct their attention to the time point indicated by the precue. The precued target and cue validity were randomly shuffled across trials, for 192 trials per precue T1 and precue T2 condition in each MEG session. Each trial was categorized as attended for the precued target and unattended for the other target, yielding 192 attended and unattended trials for each target. Note that the response cue allowed us to verify that behavior depended on cue validity, but it was irrelevant to the conditions used for decoding, as it occurred at the end of the trial. In addition, the auditory precue identity is independent of the orientation identity (vertical or horizontal), so it provides no information for decoding the stimulus representation.

The experiment also included neutral trials (20% of trials). On neutral trials, the auditory precue was a combination of the high and low tones, which directed attention to both targets and was thus uninformative. The inclusion of neutral trials allowed us to confirm the selectivity of temporal attention behaviorally (see Supplementary Figure 1and Supplementary Text). However, the neutral condition had half the number of trials as the precue T1 and precue T2 conditions and so was not included in the MEG analyses to ensure comparability across precue conditions, as decoding performance is sensitive to trial counts.

### Training

Observers first completed a behavioral training session (outside of MEG) to learn the task and determine their tilt thresholds. Tilts were thresholded individually per observer (mean tilt = 0.76°) using a 3-up-1-down staircasing procedure to achieve ∼79% accuracy on neutral trials.

### Eye tracking

Observers maintained fixation on a central circle that was 0.15° in diameter throughout each trial. Gaze position was measured using an EyeLink 1000 eye tracker (SR Research) with a sampling rate of 1000 Hz. A five-point-grid calibration was performed at the start of each session to transform gaze position into degrees of visual angle.

### MEG

Each MEG session included 12 experimental blocks that were each approximately 6 minutes long. Observers could rest between blocks and indicated their readiness for the next block with a button press.

Before MEG recording, observer head shapes were digitized using a handheld FastSCAN laser scanner (Polhemus, VT, USA). Digital markers were placed on the forehead, nasion, and the left and right tragus and peri-auricular points. These marker locations were measured at the start and end of each MEG recording session. To accurately register the marker locations relative to the MEG channels, electrodes were situated on the locations identified by the digital markers corresponding to the forehead and left and right peri-auricular points.

MEG data was continuously recorded using a 157-channel axial gradiometer (Kanazawa Institute of Technology, Kanazawa, Japan) in the KIT/NYU facility at New York University. Environmental noise was measured by three orthogonally-positioned reference magnetometers, situated roughly 20 cm away from the recording array. The magnetic fields were sampled at 1000 Hz with online DC filtering and 200 Hz high-pass filtering.

### Prepossessing

MEG preprocessing was performed in Matlab using the FieldTrip toolbox for EEG/MEG-analysis (***Oostenveld et al., 2011***) in the following steps: 1) Trials were visually inspected and manually rejected for blinks and other artifacts. The number of rejected trials per session ranged from 18 to 88 (3.49 − 17.05%), mean = 51.75 (10.03%), SD = 20.06. 2) Problematic channels were automatically identified based on the standard deviations of their recorded time series. 3) The time series from channels with extreme standard deviations were interpolated from those of neighboring channels. The number of interpolated channels per recorded session ranged from 0 to 6 (0 − 3.82%), mean = 3.85 (2.45%), SD = 1.50. 4) The time series recorded from the reference magnetometers were regressed from the channel time series to remove environmental noise.

### Peak analysis

For each session, we sorted channels by their visual responsiveness, quantified by the prominence of the evoked response peaks in the average time series across all trials. We applied the MATLAB algorithm findpeaks.m to a 300 ms window following target onset, for each target, to identify the most prominent peak per target. Peak prominence quantifies how much the peak stands out relative to other peaks based on its height and location, regardless of the directionality of the peak. Peak directionality in MEG depends on the orientation of the cortical surface with respect to the gradiometers, so visually responsive channels can show either upward or downward peaks. For each channel, we averaged peak prominence magnitude across the two targets, and ranked channels by this value. We confirmed the top ranked channels were in the posterior locations.

To assess whether temporal attention affects the evoked response amplitude and latency, we first averaged the trial time series, for each observer and precue condition, across the top *k* channels, from *k* = 1 to *k* = 50, with channels sorted by their peak prominence rankings. Channels with downward peaks were sign-flipped, so that the direction of the evoked responses was consistent across channels. To capture the early visual evoked responses in the visually responsive channels, we applied the findpeaks.m algorithm to a 100-250 ms window following each target and quantified the evoked response amplitude and latency per observer and precue condition for each channel grouping.

### Source reconstruction

To examine the cortical sources of temporal attention effects observed at the channel level, we performed source reconstruction using MNE Python (***Gramfort et al., 2013***). For each participant, a 3D mesh of the cortex was generated from their structural MRI, with an approximate resolution of 4000 vertices per hemisphere. The MEG and MRI were coregistered automatically (***Gramfort et al., 2013; Houck and Claus, 2020***) based on the three anatomical fiducial points and digitized points on the scalp scanned by the laser scanner. Forward models were computed using a single-shell Boundary Element Model (BEM), which describes the head geometry and conductivities of the different tissues. The forward model was inverted using dynamic statistical parametric mapping (dSPM) (***Dale et al., 2000***) to compute source estimates for each trial and time point. The estimated source for each vertex was a dipole that was oriented perpendicular to the cortical surface. The positive or negative value of the dipole indicated whether the currents were outgoing or ingoing, respectively (***Wang et al., 2023***). For the dSPM localization method, the typical Dipole Localization Error (DLE) is around 2 cm and Spatial Dispersion (SD) is around 4 cm (***Hauk et al., 2011; Hedrich et al., 2017***). DLE measures the Euclidean distance between the maximum of maps constructed for each dipolar source and the true source location, whereas SD quantifies the spatial spread around the true source location (***Molins et al., 2008; Hauk et al., 2011***).

We divided the brain into 34 bilateral regions defined by the Desikan–Killiany (DK) atlas (***Desikan et al., 2006***). An approximate mapping of individual ‘Desikan-Killiany’ regions of interest (ROIs) to the occipital, parietal, temporal, and frontal lobes was applied, following ***Klein and Tourville*** (***2012***).

### Decoding

We trained linear support vector machine (SVM) decoders to classify stimulus orientation (vertical vs. horizontal) at each time point (***Cichy et al., 2015; King and Dehaene, 2014***). Trials were separated into training and testing sets in a 5-fold cross-validation procedure for unbiased estimates of decoding accuracy. Separate classifiers were trained for each target, yielding a time series of decoding accuracy for each target and each precue condition. For example, when decoding T1 orientation, precue T1 trials would be attended and precue T2 trials would be unattended. To increase signal-to-noise, we averaged small numbers of trials (5 trials) to create pseudotrials (***Isik et al., 2014; Meyers, 2013; Wardle et al., 2016***) and across small time windows (5 ms) (***Isik et al., 2014***), and repeated the decoding procedure 100 times with random pseudotrial groupings to remove any idiosyncrasies due to trial averaging.

To reduce noise in the classifier, we performed feature selection in sensor space by determining the number of channels that contained the most orientation information across all trials, independent of precue condition. We compared the maximum decoding accuracy, averaged across T1 and T2, from all sessions from the most visually responsive channels, based on peak prominence (top 10, 20, 50, 100 or all channels; see *Peak analysis*) with 10 repetitions of the decoding procedure described above. The highest decoding accuracy was obtained using the top 50 channels. Therefore, for each session, we selected the top 50 visually responsive channels for sensor space decoding analysis and comparison across precue conditions. Most of the selected channels were in posterior locations. However, we note that MEG channels capture a weighted sum of the activities of all brain sources (***Pizzella et al., 2014***).

In source space, we decoded the stimulus orientation from the estimated source activation in atlas-based ROIs. Each ROI contained many vertices, whose activation time series were obtained from the source reconstruction procedure (see *Source reconstruction*). For each ROI, the number of features (vertices) can be much larger than the number of samples (trials). To avoid overfitting, we therefore reduced the feature dimension for ROIs with more than 100 vertices by univariate feature selection using ANOVA F-test (***Pedregosa et al., 2011; Gramfort et al., 2013***). ANOVA F-test feature selection was applied on the training set in the 5-fold cross-validation procedure. When training a classifier for an ROI with more than 100 vertices, we selected 100 features (i.e., estimated source activation values from 100 vertices) with the highest scores in the ANOVA F-test. Thus, the input of a classifier for a given ROI was the estimated source activation from no more than 100 vertices. To obtain the decoding performance for each of the occipital, parietal, temporal, and frontal lobes from the 34 bilateral DK ROIs, we averaged the decoding performance across the ROIs within each lobe. When investigating the left and right hemispheres separately, we decoded 68 DK ROIs with 34 DK ROIs in each hemisphere. Decoding performance in the critical time window was calculated by averaging the decoding performance across the time points in the critical time window.

### Statistical analysis

The effects of temporal attention on behavior (*d*^’^ and RT) were assessed using repeated measures ANOVAs via the pingouin package in Python. The within-subject factors were target (T1 or T2) and validity (valid or invalid, with respect to the match between the precue and the response cue), where two sessions for each subject were averaged.

The effects of temporal attention on the MEG time series (evoked response peak magnitude and latency) were assessed using repeated measures ANOVAs via the pingouin package in Python, seperately for each target and channel grouping. The within-subject factor was precue (precue T1 or precue T2), where two sessions for each subject were averaged.

To assess the effect of temporal attention on decoding performance across the full time series, we used a non-parametric test with cluster correction (***Maris and Oostenveld, 2007***). The null permutation distribution was obtained by collecting the trials of the two experimental conditions in a single set, randomly partitioning the trials into two subsets, calculating the test statistic on this random partition, and repeating the permutation procedure 1000 times to construct a histogram of the test statistic under the null hypothesis.

For each permutation, the test statistic was calculated as follows:

1. For every sample (decoding performance in a 5-ms time window), compare the decoding accuracy on the two types of trials (precue T1 versus precue T2) by means of a t-value using a paired t-test.
2. Select all samples whose t-value is larger than some threshold. Higher thresholds are better suited for identifying stronger, short-duration effects, whereas lower thresholds are better suited for identifying weaker, long-duration effects (***Maris and Oostenveld, 2007***). We selected a threshold of *t*=1.5 (*n* = 10 subjects), where two sessions for each subject were averaged.
3. Cluster the selected samples in connected sets on the basis of temporal adjacency.
4. Calculate cluster-level statistics by taking the sum of the t-values within a cluster.
5. Take the largest of the cluster-level statistics.

The spatial cluster permutation for Figure 4A was calculated in a way similar to the steps described above using the MNE package in Python with permutation_cluster_1samp_test function, where the adjacency matrix for the function was determined based on the anatomical surface location of the DK ROIs, and the number of permutations n_permutations was set to “all” to perform an exact test. For each ROI, the averaged decoding accuracy across the time points in the critical time window for the two types (precue T1 versus precue T2) of trials were compared using a paired t-test with threshold *t*=2.1 (*n* = 20 sessions), alpha level 0.05.

## Acknowledgments

We thank Sirui Liu and Luis Ramirez for assistance with early versions of the experiment. We thank Hsin-Hung Li as well as members of the Denison and Carrasco Labs for helpful comments. We thank Jeffrey Walker at the NYU MEG lab for technical assistance. This research was supported by National Institutes of Health National Eye Institute R01 EY019693 to M.C., F32 EY025533 to R.N.D., National Defense Science and Engineering Graduate Fellowship to K.J.T., T32 EY007136 to NYU, and by funding from Boston University to R.N.D.

## Data availability

The data underlying this article are available in Open Science Framework (OSF) at https://osf.io/7u6x3/, and can be accessed with DOI 10.17605/OSF.IO/7U6X3.

## Supplementary Text

### Temporal attentional selection in two-target temporal cueing task

Perceptual sensitivity was impaired in the neutral condition (when the temporal precue was uninformative) compared to the valid condition for T1 (see Supplementary Figure 1), confirming that observers could not fully process both stimuli and instead used attention to select the more relevant stimulus in the sequence. Such behavioral benefits are consistent with previous studies that used the two-target temporal cueing task (***Denison et al., 2017, 2021; Fernández et al., 2019; Duyar et al., 2024***), confirming that this task reliably elicits temporal attentional selection.

### Replication of enhanced T1 orientation decoding with temporal attention

We confirmed the enhancement of temporal attention on orientation decoding for T1 using an identical analysis procedure in a separate dataset, in which the targets were superimposed on a 20-Hz flickering noise patch instead of a blank background (see Supplementary Figure 2A). Although this experiment was not designed for decoding analysis due to the continuous presence of flickering noise, we again found an enhancement of orientation representation in attended vs. unattended trials at a similar time window around 250 ms (195-260 ms) after target onset (Supplementary Figure 2B). Again, there was no effect of temporal attention on T2 decoding performance. The overlap of the time windows in which temporal attention enhanced orientation representations in the two experiments (235-260 ms after target onset) indicates that temporal attention reliably affects the orientation representation in an intermediate processing time window following the earliest visual evoked responses and peak decoding accuracy.

### Brief early peak in decoding accuracy for T1

Although only the “critical window” around 250 ms passed the stringent cluster correction test across the full trial time series, we noted a brief early peak (at 90 ms after target onset, uncorrected p = 0.019) in decoding accuracy for T1 that appeared to be present when T1 was attended but absent when it was unattended (See Supplementary Figure 3A). Given a previous finding that temporal attention transiently affects evoked responses to steady-state visual stimulation (***Denison et al., 2024***), we used source reconstruction to investigate the cortical origin of this early peak modulation. The effects of temporal attention at 90 ms were strongest in occipital and parietal areas (Supplementary Figure 3B), a strikingly different topography from the fronto-cingulate areas modulated during the later critical time window. This result suggests that any effect of temporal attention on early stimulus representations is localized to visual areas.

## Supplementary Figures

**Supplementary Figure 1.**
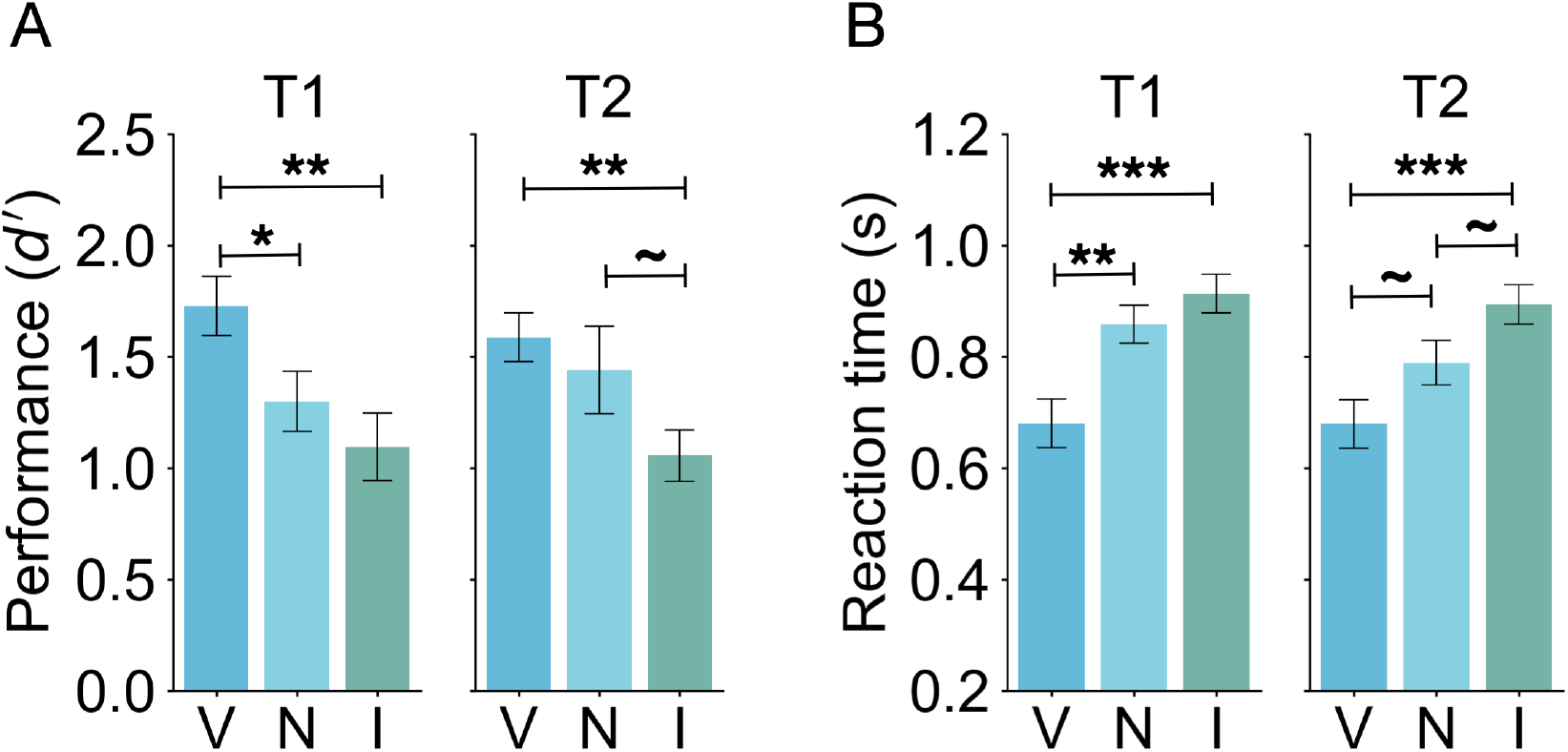
Behavioral results including the neutral condition. (A) Tilt discrimination (sensitivity) and (B) reaction time for each target (T1, T2) and validity condition. Error bars indicate ±1 SEM. ∼ p *<* 0.1, * p *<* 0.05, ** p *<* 0.01; *** p *<* 0.001.

**Supplementary Figure 2.**
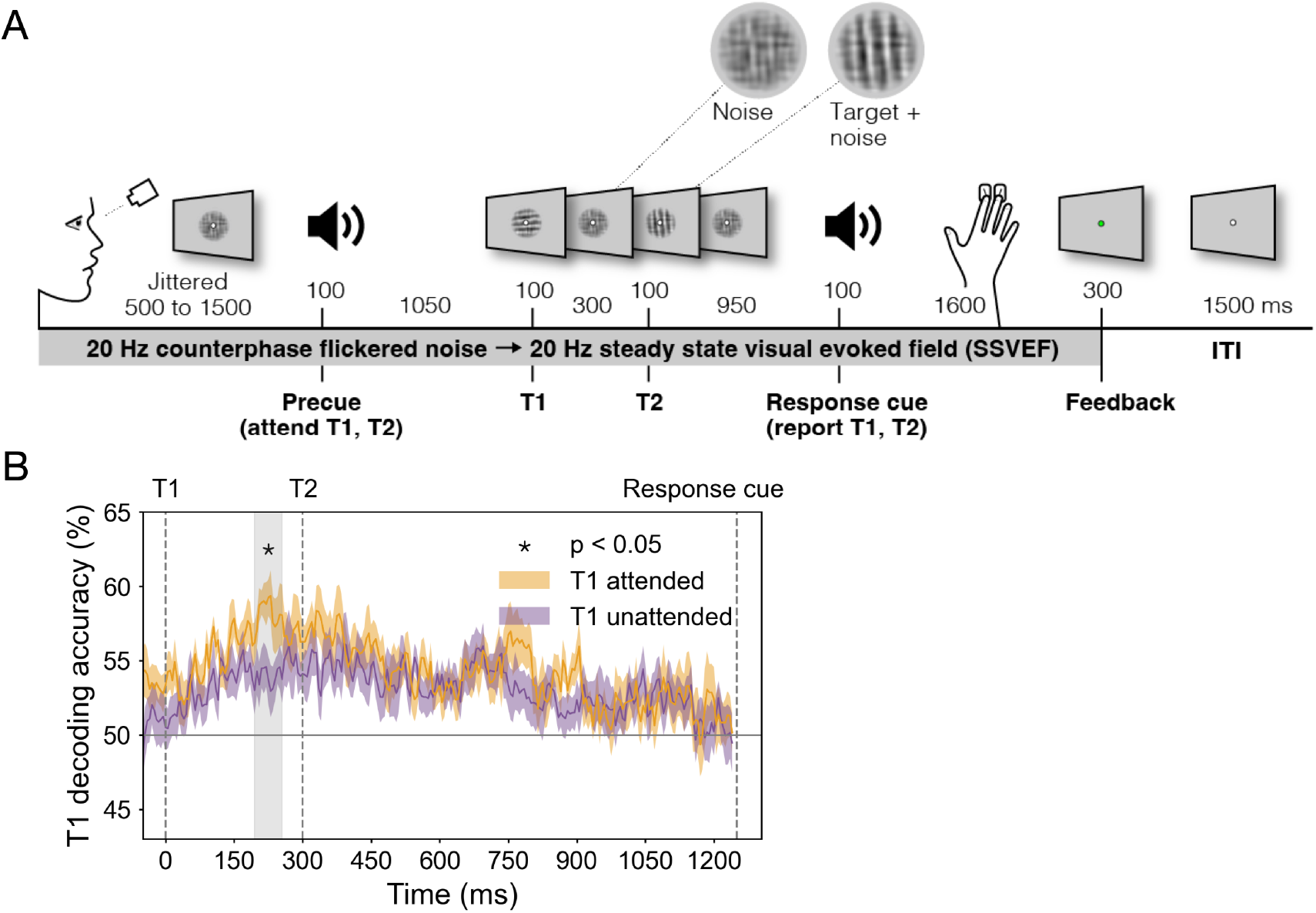
Decoding performance for a separate experiment with targets superimposed on flickering noise. (A) Two-target temporal cueing task. Trial timeline showing stimulus durations and SOAs. Targets were embedded in 20 Hz counterphase flickering noise. Precues and response cues were pure tones (high = cue T1, low = cue T2). (B) T1 orientation decoding performance for T1 attended and T1 unattended trials confirms enhancement of T1 representation in an intermediate time window.

**Supplementary Figure 3.**
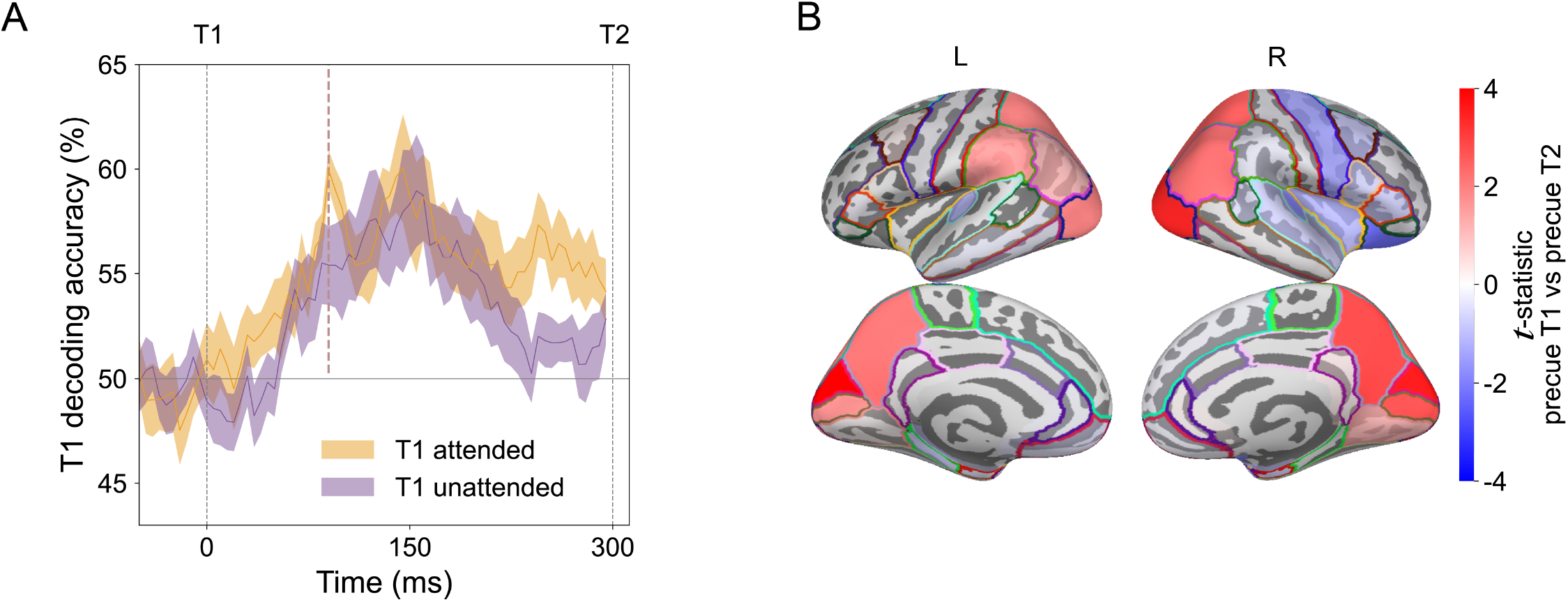
Topography of attentional enhancement of orientation representations during an early peak. (A) T1 decoding time series for attended and unattended trials highlighting early peak (thick dashed line, same data as in Figure 3C). (B) T1 decoding differences between attended and unattended conditions for left (L) and right (R) hemispheres at 90 ms after target onset for each of 68 DK ROIs.

